# Structural and biochemical studies of an NB-ARC domain from a plant NLR immune receptor

**DOI:** 10.1101/557280

**Authors:** John FC Steele, Richard K Hughes, Mark J Banfield

## Abstract

Plant NLRs are modular immune receptors that trigger rapid cell death in response to attempted infection by pathogens. A highly conserved <u>n</u>ucleotide-<u>b</u>inding domain shared with <u>A</u>PAF-1, various <u>R</u>-proteins and <u>C</u>ED-4 (NB-ARC domain) is proposed to act as a molecular switch, cycling between ADP (repressed) and ATP (active) bound forms. Studies of plant NLR NB-ARC domains have revealed functional similarities to mammalian homologues, and provided insight into potential mechanisms of regulation. However, further advances have been limited by difficulties in obtaining sufficient yields of protein suitable for structural and biochemical techniques. From protein expression screens in *Escherichia coli* and *Sf9* insect cells, we defined suitable conditions to produce the NB-ARC domain from the tomato NLR NRC1. Biophysical analyses of this domain showed it is a folded, soluble protein. Structural studies revealed the NRC1 NB-ARC domain had co-purified with ADP, and confirmed predicted structural similarities between plant NLR NB-ARC domains and their mammalian homologues.

## Introduction

Plants mount sophisticated responses to attack by pathogens following the detection of specific microbe-derived signals. General responses including callose deposition (1), production of reactive oxygen species (2), and upregulation of defence-specific genes (3) can be triggered by the detection of conserved microbe-derived pathogen-associated molecular patterns (PAMPs) such as fungal chitin (4, 5) or bacterial flagellin (6, 7) at the cell surface.

Such responses are generally effective at preventing or reducing colonisation of plants by non-adapted microbes, however they are less effective against specialised plant pathogens. Various sub-populations of plant pathogens have evolved to infect specific cultivars of a given host plant in a gene-for-gene manner (8, 9). A key feature of biotrophic or hemi-biotrophic plant pathogens are translocated effector proteins that enter the host cell, typically to subvert cell surface-based immune responses (10-13). Plants have in turn evolved intracellular receptors capable of detecting the presence (14-16) or activity (17, 18) of such effectors. Upon detection, these receptors trigger an immune response, typically characterised by localised programmed cell-death known as the hypersensitive response (19).

The majority of these plant intracellular receptors belong to the nucleotide-binding leucine-rich repeat (NLR) protein family (20, 21), similar to those involved in certain animal cell-death or innate immune responses (22-24). Plant NLRs share a common architecture with animal NLRs, containing nucleotide binding (NB) and Leucine-rich repeat (LRR) domains, but can be further classified into CNLs and TNLs, dependent on whether they have an N-terminal coiled-coil (CC) domain or N-terminal toll/interleukin-1 receptor (TIR) domain. Recently, variations on this general structural framework have been identified that include the integration of additional domains into the classical NLR architecture (25, 26).

Bioinformatic and *in planta* studies have assigned specific functions to plant NLR domains. The N-terminal CC or TIR domains are typically described as required for downstream signalling following perception of pathogens (27-30), the LRR is implicated in auto-inhibition and/or effector detection (31-34), and the central NB-ARC domain is posited as a regulatory domain (35, 36), acting as a “switch” that determines whether the protein is in an active or inactive state (37). In plant NLRs, this central NB-ARC domain is presumed to be structurally related to a large family of nucleotide binding proteins (38), and biochemical investigations using protein refolded from *E. coli* inclusion bodies indicate that plant NB-ARC domains are able to bind to, and hydrolyse, ATP (39). Expression of full-length plant NLRs such as MLA27 (29) and the flax rust resistance protein M (40) have shown that full-length proteins co-purify with ADP, similar to what has been observed for mammalian NLRs (41-43). Further investigation of full-length plant NLRs have been hampered by low yields in expression systems, and more sensitive biochemical and structural investigations addressing the NB-ARC domain are constrained by the use of refolded (39), or truncated protein (44). This leaves many fundamental explanations for mutant phenotypes reliant on predicted similarities to mammalian homologues, notably the well-characterised APAF-1 protein (37, 45-48).

NB-ARC domains of animal (also known as NACHT domains) and plant NLRs belong to the AAA+ protein superfamily, with several shared motifs known to contribute to protein function. Key amongst these are the Walker A motif (or P-loop), which is important for nucleotide binding, and Walker B motif, which is required for ATP hydrolysis. Well-established mutations that target these sites include replacing the charged lysine residue in the Walker A motif with alanine or arginine, resulting in a loss of nucleotide binding, and mutating the charged amino acids of the Walker-B ΦΦΦΦD(D/E) motif (four hydrophobic amino acids followed by an aspartate then either a second aspartate or glutamate) to alanine or glutamine, which inhibits ATPase activity (84, and references therein). Additional NB-ARC motifs that have been studied include the conserved “GLPL” (glycine-leucine-proline-leucine) motif, and the “MHD” motif (methionine-histidine-aspartate) which, when mutated, usually results in an autoactive phenotype (34, 53). Biochemical investigation of the flax resistance protein M was able to link autoactivation caused by disruption of the MHD motif to the state of the bound nucleotide (40).

Here we characterise the NB-ARC domain from the tomato NLR NRC1, which we have expressed and purified from both bacterial (*E. coli*) and insect (*Sf9*) cell culture. Biophysical analysis of the NRC1 NB-ARC domain by circular dichroism, analytical gel filtration, and differential scanning fluorimetry indicate the protein is a well-folded, stable monomer in solution. We also crystallised and determined a structure of the NRC1 NB-ARC domain. The structural analysis highlights the protein’s similarity to the equivalent region of the mammalian NLR APAF-1, and identified co-purifying ligands.

## Results

### Defining the boundaries of the NRC1 NB-ARC domain

We used several bioinformatics resources to help define the boundaries of the NB-ARC domain from the canonical CNL NRC1 (49), with the aim of producing soluble, stable protein (Figure 1). A putative NB-ARC domain spanning residues 165 to 441 was identified by Pfam (50, 51). However, this region did not include the conserved MHD motif (valine-histidine-aspartate, VHD, in NRC1) towards the C-terminus of the domain, a region known to be functionally relevant (34, 47, 49, 52, 53). Therefore, we extended our analyses to include a combination of LRR-specific bioinformatic searches (LRR Finder (54) and LRRsearch (55)), secondary structure prediction (Phyre2 (56)), and disorder predictions (RONN (57)) to define a C-terminal boundary of the NRC1 NB-ARC domain. Based on the results of these searches, we assigned the NRC1 NB-ARC domain C-terminus at residue 494. The secondary structure predictions also suggested that truncating the protein at position 165 (putative N-terminus) could disrupt an α-helix. With this information, we re-assigned the NRC1 NB-ARC domain N-terminus at position 150, within a predicted disordered region between the CC and NB-ARC domains.

**Figure 1.**
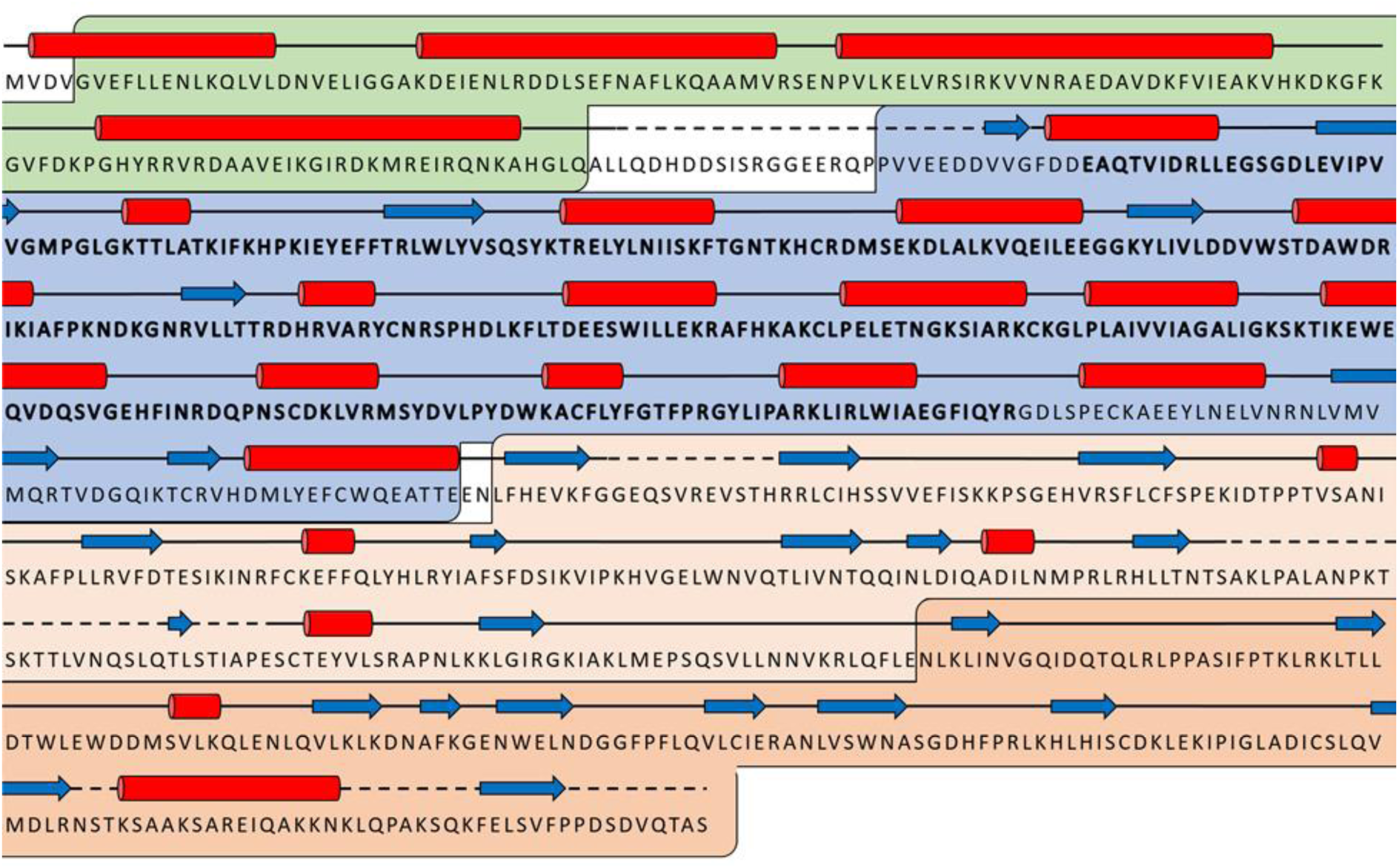
NRC1 domain assignment. The extent of the coiled-coil domain was predicted as residues 4-134 (green) by CCHMM (71) and MARCOILS (72), the NB-ARC domain to be residues 150 to 494 (blue) by Pfam (50, 51) (predicted domain spanning residues 165-441 in bold), and the LRR domain from 718-888 (dark orange) by LRR Finder (54) and LRRsearch (55). Light orange indicates a putative LRR region. Predicted secondary structure elements are shown as red cylinders (helices) and blue arrows (beta-strands), as defined by Phyre2 (56), and dashed lines (disordered loop regions) from RONN (57).

### Expression and purification of wild-type NRC1 NB-ARC domain

Based on our bioinformatic analysis, a construct spanning NRC1 residues 150 to 494 was cloned into both the pOPIN-F and pOPIN-S3C expression vectors (58), which allow for protein expression with a cleavable N-terminal 6xHis tag or N-terminal 6xHis-SUMO (small ubiquitin-like modifier protein) tag respectively.

We used the high-throughput capacity of Oxford Protein Production Facility, UK, to screen for expression/purification of the NRC1 NB-ARC domain in various *E. coli* strains/media and in insect cells. We found the best conditions for soluble expression used pOPIN-S3C in *E. coli* strain Lemo21(DE3), grown in PowerBroth ™ media, and baculovirus-mediated expression in *Sf9* cells using a pOPIN-F construct. Using a combination of immobilised metal ion chromatography (IMAC) followed by size-exclusion chromatography (SEC) we were able to purify a protein of molecular weight consistent with the 6xHis SUMO-tagged NRC1 NB-ARC domain from *E. coli* lysates (Figure 2a). Removal of the N-terminal tag (with 3C protease) followed by subtractive IMAC and a second round of SEC resulted in protein at high purity (Figure 2b). A similar purification procedure for 6xHis-tagged protein expressed in *Sf9* cells also yielded pure NRC1 NB-ARC protein (Figure 2b). Intact mass spectrometry confirmed the masses of the proteins from both expression systems were as expected.

**Figure 2.**
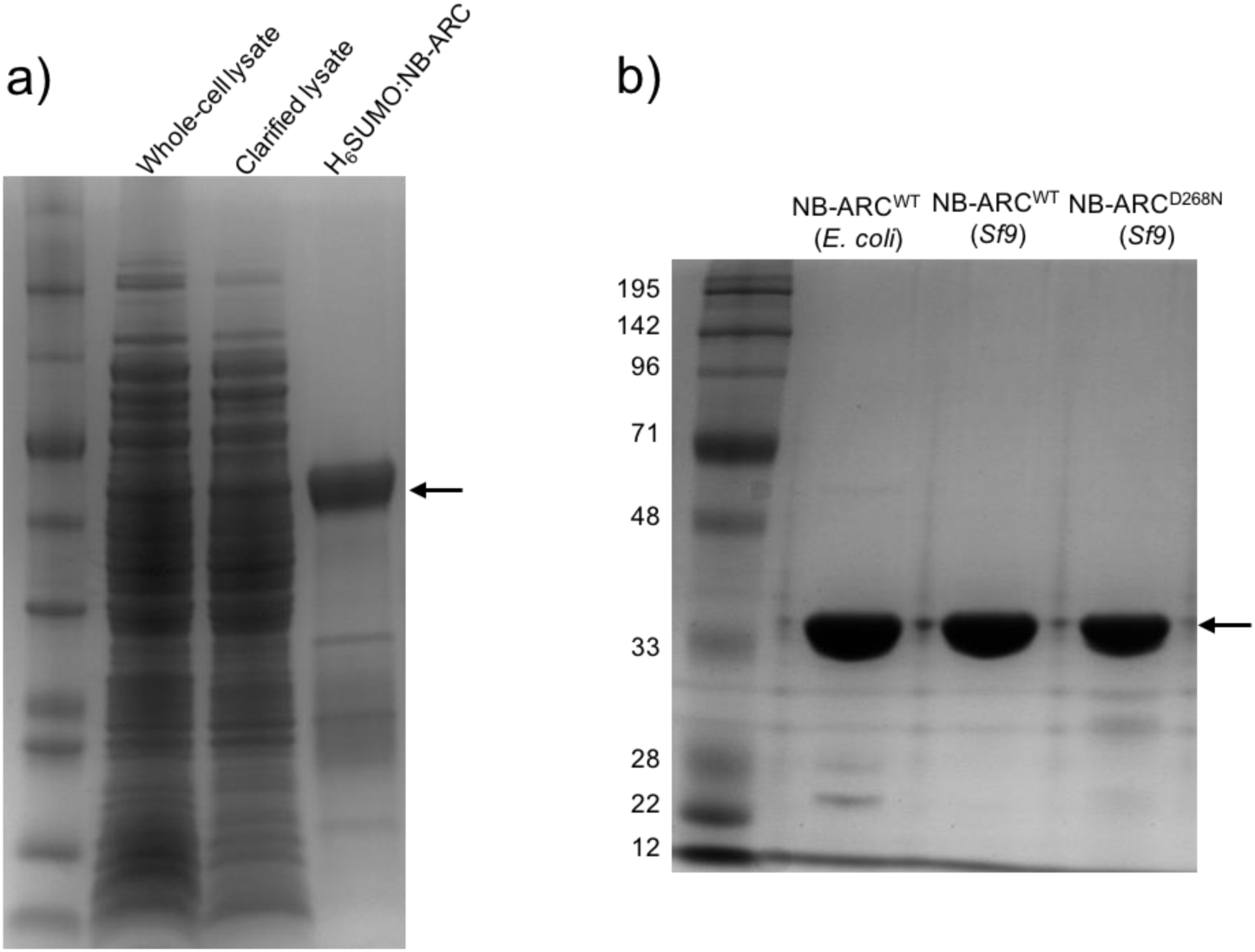
Purification of the NRC1 NB-ARC domain from *E. coli* and *Sf9* cells. a) SDS-PAGE showing the purification of NRC1 NB-ARC from *E coli* lysates. Lane 1. Molecular weight marker, 2. Whole-cell lysate, 3. Total soluble protein, 4. 6xHis-SUMO tagged NRC1 NB-ARC following one-step IMAC/gel filtration purification. b) SDS-PAGE showing final purified protein following N-terminal tag removal. Lane 1. Molecular weight marker, 2. Wild-type protein purified from *E. coli*, 3. Wild-type protein purified from *Sf9* cells, 4. NRC1 NB-ARC^D268N^purified from *Sf9* cells. Black arrow indicates position of NRC1 NB-ARC protein.

### Expression and purification of NRC1 NB-ARC domain mutants

We designed several point mutants in the NRC1 NB-ARC domain to perturb sequence motifs with established roles in animal and plant NLRs. These included regions important for ATP-binding (Walker-A motif; K191A), ATP hydrolysis (Walker-B motif; D268N, D267N, D267N/D268N) and NLR regulation (MHD motif; D481V, H480F, H480A). Unfortunately, we were unable to express and purify any of these mutants in *E. coli*, despite screening multiple strains (including BL21(DE3), BL21*, soluBL21, Rosetta2 pLysS, Lemo21(DE3), and BL21-AIpLysS) at either 37°C or 18°C with different growth media (LB, Terrific Broth, Powerbroth), or with additives to the media to improve soluble protein expression (DMSO, glycerol (59), or ethanol (60)).

Having identified that baculovirus-mediated expression in *Sf9* cells was able to generate soluble wild-type protein, we returned to insect-cell expression to screen NB-ARC mutant constructs using the pOPIN-F vector. With these screens, we could produce the mutant D268N (Walker B motif), and purify it to homogeneity in yields similar to the wild-type protein (Figure 2b). The other mutants were recalcitrant to soluble expression/purification. Intact mass spectrometry confirmed mass for the D268N mutant was as expected.

### Biophysical characterisation of the NRC1 NB-ARC domain

To investigate the biophysical properties of wild-type NRC1 NB-ARC domain, and the NB-ARC^D268N^mutant, we used three approaches: analytical gel filtration, circular dichroism, and thermal denaturation.

Firstly, we used analytical gel filtration and showed that both wild-type protein (produced in *E. coli* or insect cells) and the D268N mutant behave as monomers in solution, and elute at a retention volume consistent with their molecular mass (Figure 3a,b).

**Figure 3.**
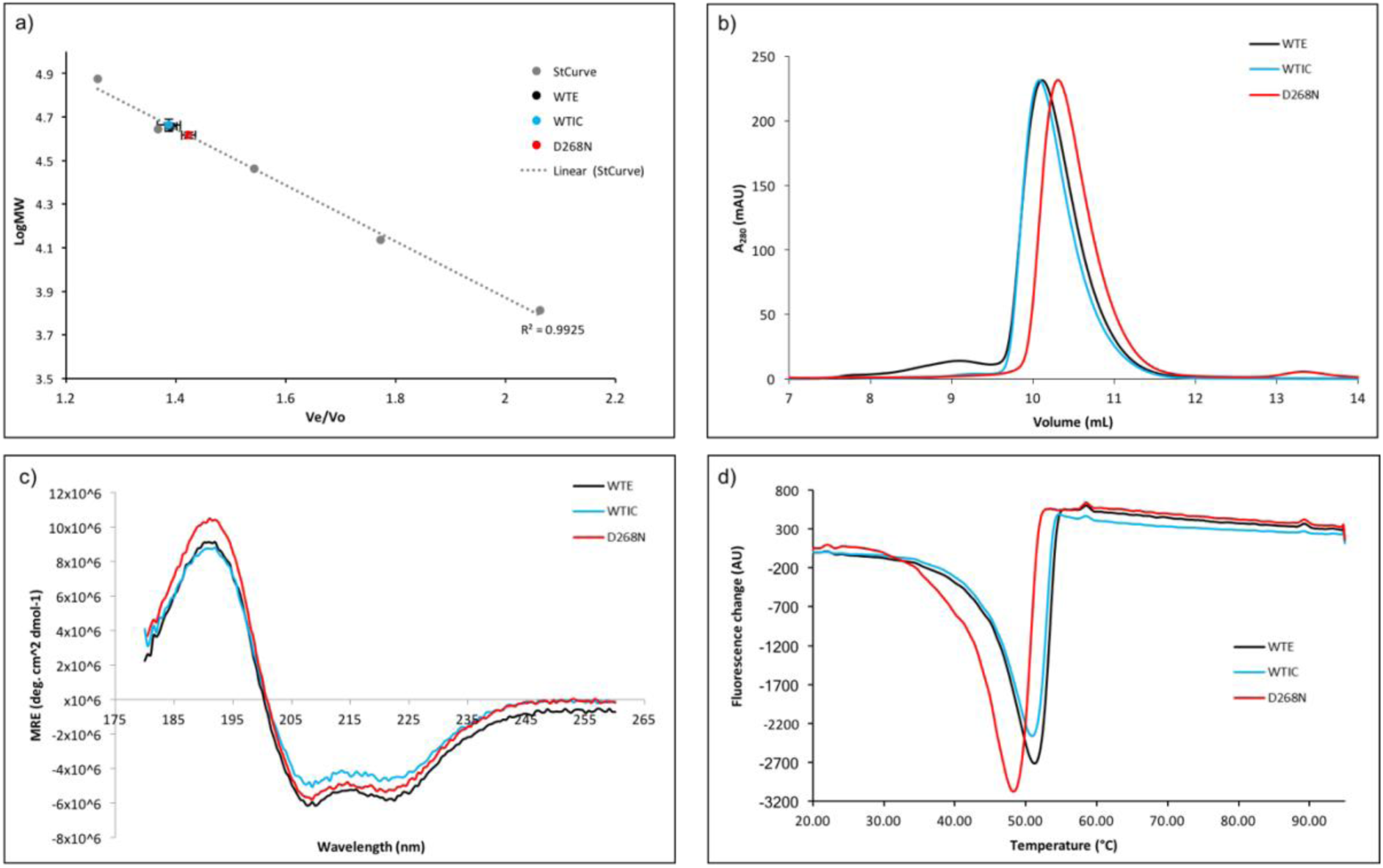
Biophysical characterisation of NRC1 NB-ARC proteins. a) Elution of NRC1 NB-ARC proteins plotted on a standard curve determined from the elution of Aprotinin (6.5kDa), Ribonuclease A (13.7kDa), Carbonic anhydrase (29.0kDa), Ovalbumin (44.0kDa), and Conalbumin (75.0kDa), with dextran used to calculate column void volume. Error bars represent standard error of the mean from three experiments using two independent purifications. b) Elution of purified NRC1 NB-ARC domain proteins by analytical gel filtration. Single elution peaks were observed for wild-type proteins at ∼10.2mL, with the D268N mutant eluting slightly later, at ∼10.5mL. c) Circular dichroism spectra of wild-type and mutant NRC1 NB-ARC domains is consistent with folded protein. d) First derivative plot of fluorescence against temperature during thermal melting differential scanning fluorimetry experiments of wild-type and mutant NRC1 NB-ARC proteins. WT-E: Wild-type NRC1 NB-ARC domain expressed in *E. coli*, WT-IC: Wild-type NRC1 NB-ARC domain expressed in *Sf9* cells, D268N: mutated NRC1 NB-ARC domain.

Secondly, we assessed NRC1 NB-ARC domain folding by circular dichroism (CD). Each wild-type sample gave a spectrum consistent with a well-folded protein, possessing mixed secondary structure (Figure 3c, Table 1). Disruption of the Walker B motif (with the NRC1 NB-ARC^D268N^mutant) had little effect on protein secondary structure as measured by CD (Figure 3c, Table 1).

**Table 1.**
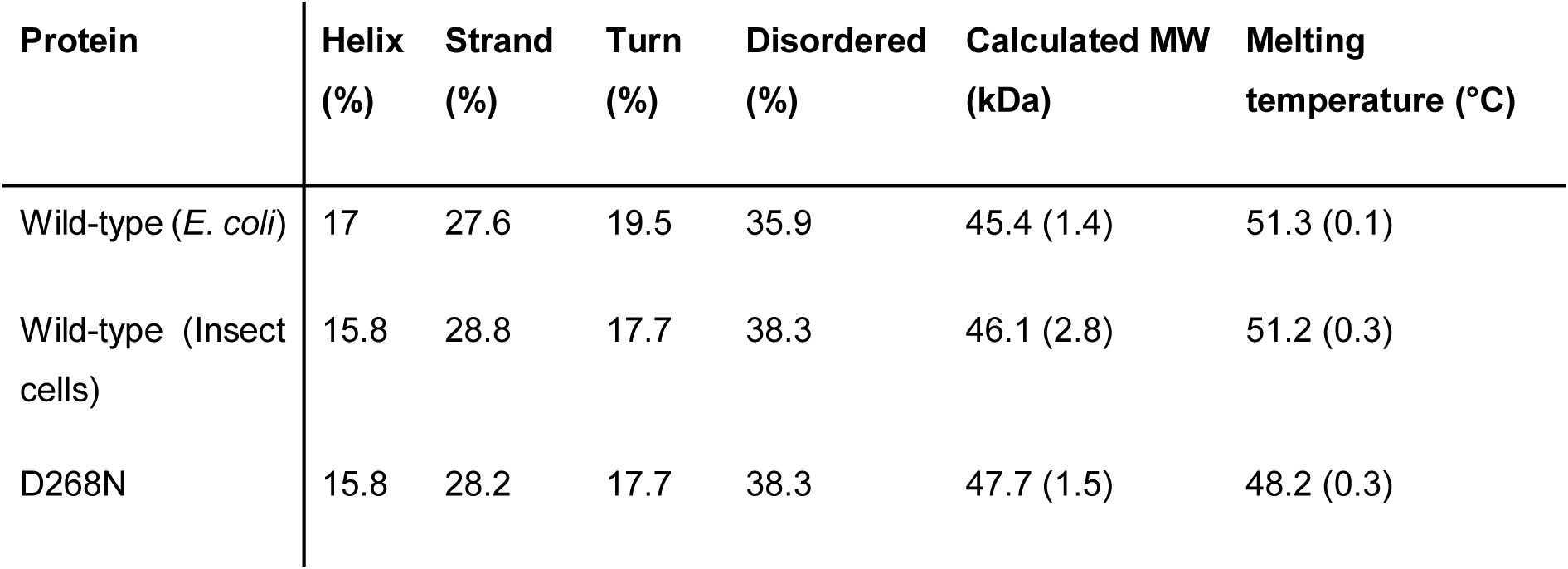
Summary of the biophysical properties of NRC1 NBARC domains. Secondary structure elements expressed as % of protein fold were calculated using Contin on the Dichroweb server (82,83) from CD data. NRMSD for assignments are 0.103, 0.143 and 0.099 for WT-*E. coli*, WT-insect cells and D268N mutant data respectively. Calculated molecular weights are from gel filtration, and melting temperatures from DSF. Values in parentheses represent standard error for calculated molecular weights and standard deviation of the mean for melting temperatures.

| Protein | Helix (%) | Strand (%) | Turn (%) | Disordered (%) | Calculated MW (kDa) | Melting temperature (°C) |
| --- | --- | --- | --- | --- | --- | --- |
| Wild-type (*E. coli*) | 17 | 27.6 | 19.5 | 35.9 | 45.4 (1.4) | 51.3 (0.1) |
| Wild-type (Insect cells) | 15.8 | 28.8 | 17.7 | 38.3 | 46.1 (2.8) | 51.2 (0.3) |
| D268N | 15.8 | 28.2 | 17.7 | 38.3 | 47.7 (1.5) | 48.2 (0.3) |

Finally, we used differential scanning fluorimetry (DSF) to assess the thermal stability of the three proteins. In this assay, an increase in Sypro orange(™) fluorescence is associated with a transition from a folded protein to denatured state as internal hydrophobic surfaces are exposed. For all three NB-ARC proteins, a single inflection point for change in fluorescence was observed at ∼50°C, consistent with a single unfolding event (Figure 3d, Table 1).

Taken together, these data are consistent with the wild-type NRC1 NB-ARC domain, and D268N mutant, adopting a stable, folded state in solution.

### NRC1 NB-ARC domain purifies in an ADP-bound form

The full length flax NLR protein M, and the barley NLR MLA27, were both shown to co-purify with ADP as a bound nucleotide, with a small fraction of MLA27 bound to ATP (29, 40). Individual NB-ARC domains from animal systems have also been shown to co-purify with nucleotides, but these vary between ADP and ATP bound states (42, 61, 62). We employed a luciferase-based assay (40, also as described in the Materials and Methods) to quantify the relative occupancy of these nucleotides in the purified NRC1 NB-ARC domain. Using this assay, we were unable to measure any ATP co-purifying with the protein. We then converted any ADP present to ATP using pyruvate kinase, and used this as a proxy for bound ADP. As no ATP was bound to the protein, we express ADP occupancy as a percentage, calculated from relative protein/nucleotide concentrations. These data show that all three NRC1 NB-ARC proteins co-purified with ADP with high occupancy (Figure 4). Based on the higher affinity of APAF-1 for ATP over ADP (70), and ATPase activity displayed by other NB-ARC proteins, we expect this high ADP occupancy is due to intrinsic ATPase activity of NRC1 NB-ARC domain. Surprisingly, NRC1 NB-ARC^D268N^, a mutant designed to abolish ATPase activity, also co-purified with ADP suggesting either preferential binding of ADP over ATP, or residual ATPase activity.

**Figure 4.**
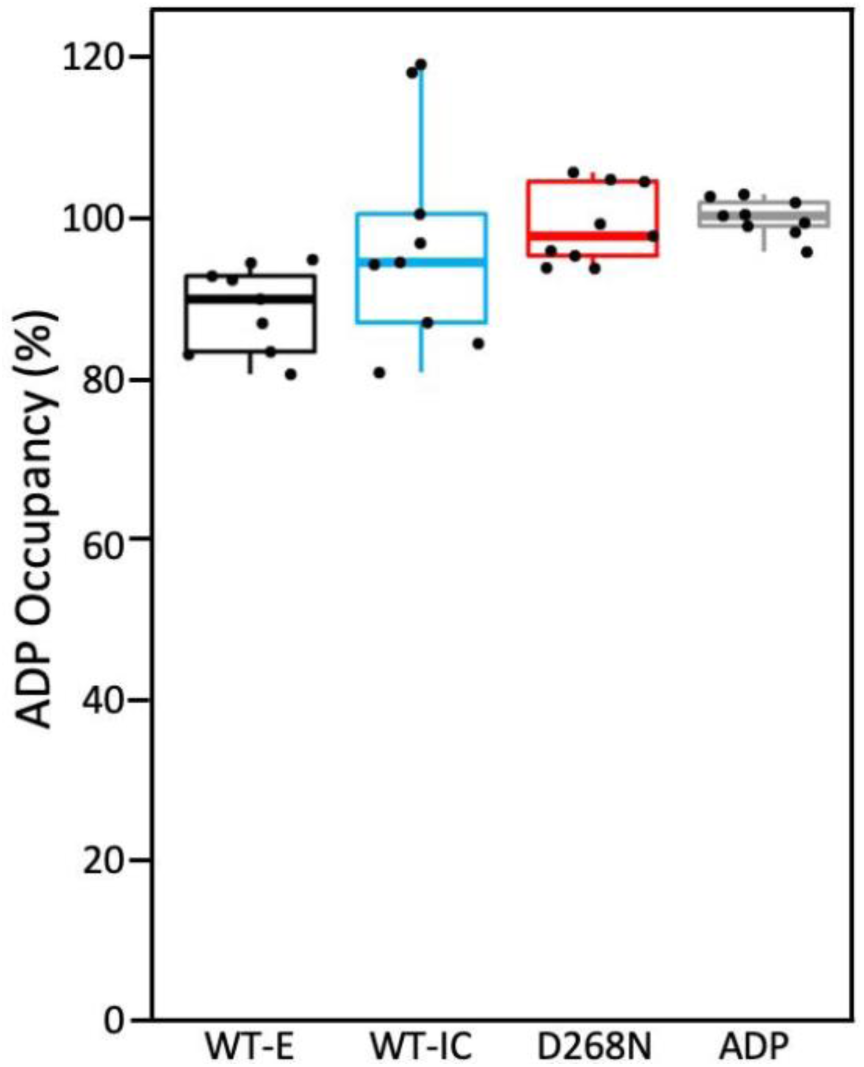
Quantification of nucleotide co-purifying with NRC1 NB-ARC proteins. As assays were unable to detect bound ATP, a percentage occupancy of each protein with ADP is shown. Experiments were performed in triplicate with three technical replicates. ADP was included as an internal standard for each run. The centre line represents the median, the box top and bottom limits are the upper and lower quartiles, respectively, and the whiskers are the 1.5×interquartile range. All of the data points are represented as dots, with no outliers present. WT-E: Wild-type NRC1 NB-ARC domain expressed in *E. coli*, WT-IC: Wild-type NRC1 NB-ARC domain expressed in *Sf9* cells, D268N: mutated NRC1 NB-ARC domain.

### NRC1 NB-ARC domain shows structural similarities to APAF-1

We generated a selenomethionine (SeMet) derivative of the NRC1 NB-ARC domain, which readily crystallised to form long, tapered rods. Crystals were flash frozen in liquid nitrogen and X-ray data collected at the Diamond Light Source (UK). The resulting data were processed to 3.03 Å resolution. Isomorphous native crystals of the NRC1 NB-ARC domain were generated, and X-ray data collected to 2.5 Å resolution (Table 2).

**Table 2.** X-ray data collection and refinement statistics for the native and SeMet NRC1 NB-ARC domain.

|  | NRC1 NBARC<br>Selenomethionine | NRC1 NBARC<br>Native |
| --- | --- | --- |
| **Data collection statistics** |  |  |
| Wavelength (Å) | 0.9797 | 0.9795 |
| Space group | P6<sub>4</sub> | P6<sub>4</sub> |
| Cell dimensions |  |  |
| *a*, *b*, *c* (Å) | 110.50, 110.50, 53.55 | 110.52, 110.52, 53.34 |
| Resolution (Å)* | 38.45-3.03 (3.08-3.03) | 50.00-2.50 (2.60-2.50) |
| *R*<sub>merge</sub> (%) | 11.0 (217.7) | 5.4 (73.6) |
| *I*/*σI* | 23.6 (1.9) | 24.0 (3.5) |
| Completeness (%) |  |  |
| Overall | 100 (100) | 100 (100) |
| Anomalous | 100 (5.1) |  |
| Unique reflections | 7415 (372) | 13066 (1463) |
| Redundancy |  |  |
| Overall | 36.8 (25.2) | 10.3 (10.3) |
| Anomalous | 19.2 (13.0) |  |
| CC(1/2) (%) | 100 (71.0) | 1.00 (74.5) |
| **Refinement and model statistics** |  |  |
| Resolution (Å)* |  | 50.00-2.50 (2.60-2.50) |
| *R*<sub>work</sub>/*R*<sub>free</sub> (%) |  | 30.5 (40.1)/33.7 (45.0) |
| No. atoms |  | 2081 |
| Protein |  | 2054 |
| Ligand |  | 27 |
| B-factors |  |  |
| Protein |  | 83.6 |
| Ligand |  | 62.4 |
| R.m.s deviations |  |  |
| Bond lengths (Å) |  | 0.002 |
| Bond angles (°) |  | 0.487 |
| Ramachandran plot (%)** |  |  |
| Favoured |  | 92.6 |
| Outliers |  | 1.3 |
| MolProbity Score |  | 1.71 (99<sup>th</sup> percentile) |

\*The highest resolution shell is shown in parentheses.
\*\*As calculated by MolProbity

Structure solution (by single wavelength anomalous dispersion (SAD)) and auto-building using the Phenix autosol pipeline, with subsequent phase extension to the high-resolution dataset, yielded an interpretable electron density map. The initial autobuilding placed 171 residues in 15 fragments. Iterative manual rebuilding and refinement resulted in a final structure consisting of 252 of a total 346 residues present in the expressed protein construct. Several regions of the protein could not be modelled due to discontinuous electron density, including much of the ARC1 subdomain, and the solvent-exposed ?-helices of the Rossmann fold. Submission of this model to the DALI server (63) returns APAF-1 as the major structural homologue, despite sharing only 24% sequence identity. Side-by-side comparison of the NRC1 NB-ARC domain structure with the corresponding region from APAF-1 highlights the overall structural similarity in subdomain organisation (Figure 5).

**Figure 5.**
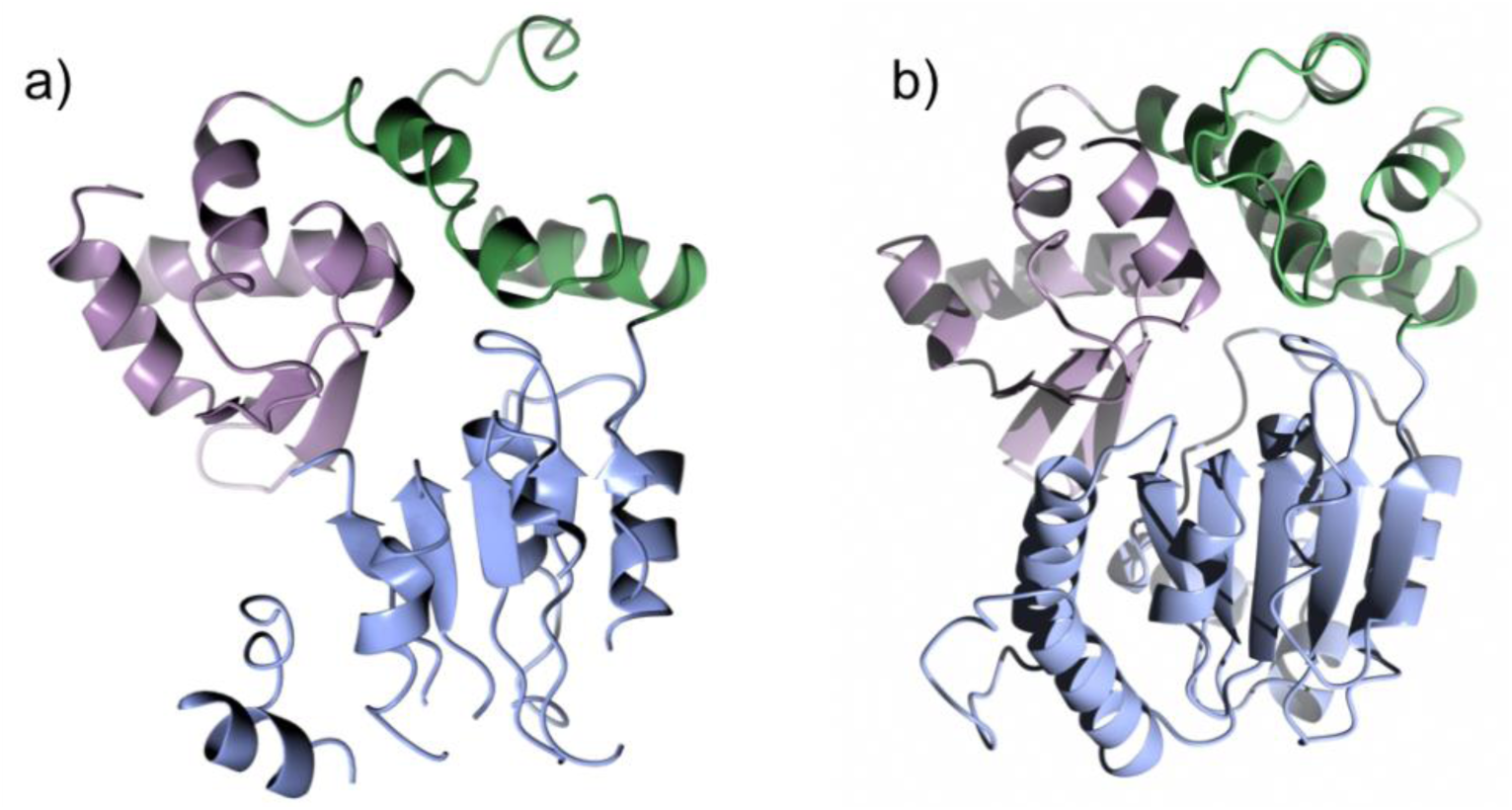
Structure of the NRC1 NB-ARC domain compared to APAF-1. Domain organisation of the NRC1 NB-ARC domain (a, left) and the equivalent region of APAF-1 (b, right, PDB: 1z6t). Subdomains are colour-coded showing the nucleotide binding (NB) region in blue, ARC1 in green, and ARC2 in lilac.

### ADP is bound in the nucleotide binding pocket of NRC1 NB-ARC domain

The electron density within the nucleotide-binding pocket of the NRC1 NB-ARC domain, and the surrounding regions, was well resolved and allowed us to unambiguously position a molecule of ADP, and the side-chains of interacting residues. These included lysine 191 of the Walker A (P-loop) motif, aspartate 267 and 268 of the Walker B motifs, and histidine 480 of the MHD motif (Figure 6a).

**Figure 6.**
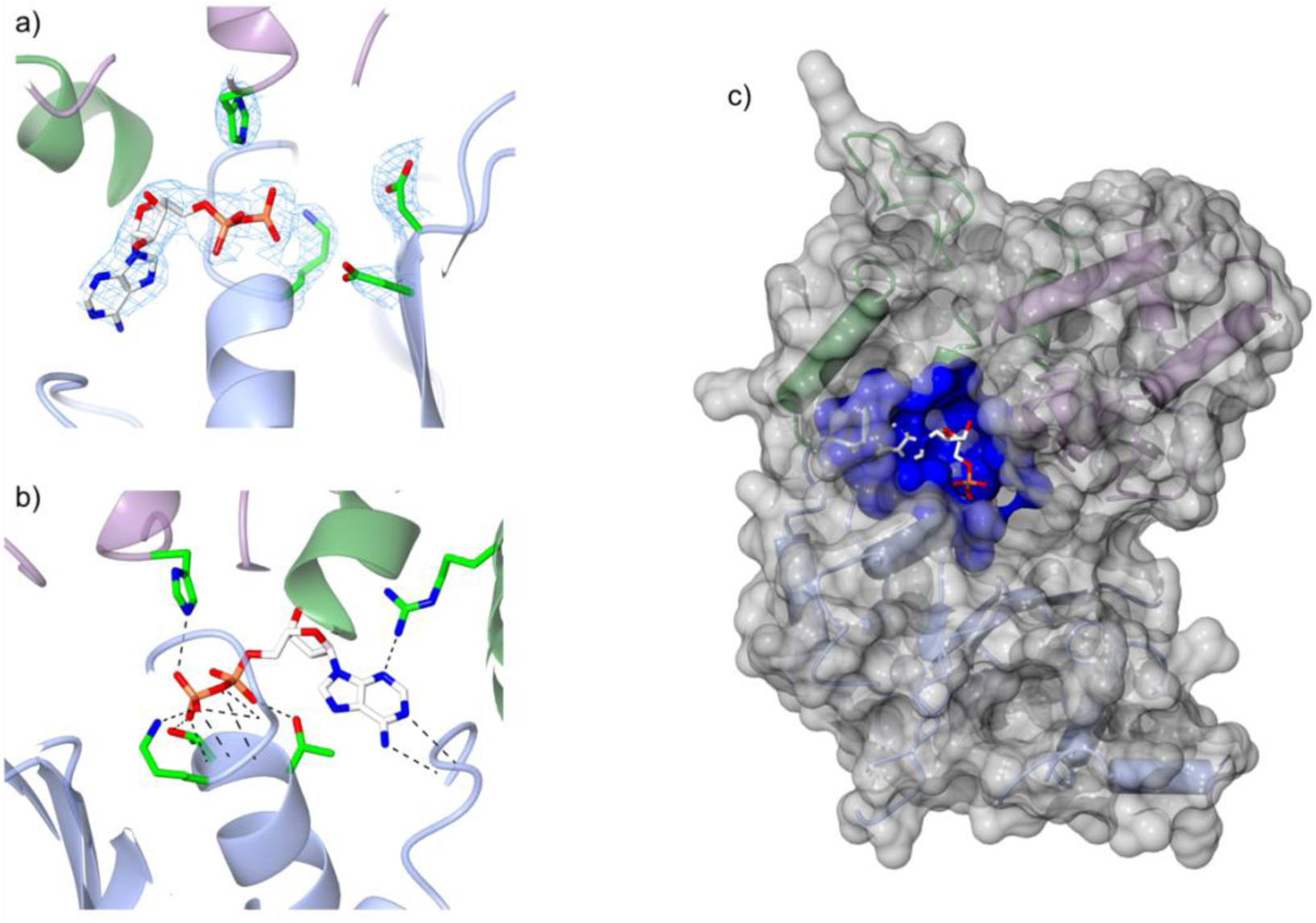
The NRC1 NB-ARC domain binds ADP. a) The ligand binding site of the NRC1 NB-ARC domain showing ADP modelled in the electron density from the final refinement. Also shown are histidine 480 of the MHD motif, lysine 191 of the Walker A motif, and aspartate 267 and aspartate 268 of the Walker B motif. b) Dashed lines depicting the interactions between protein and bound ADP, as identified by PDBe PISA (64). c) Surface representation of the NRC1 NB-ARC domain, with regions forming contacts with ADP highlighted in dark blue. The NB-ARC sub-domains are coloured as in Figure 5.

Using PDBe PISA (64), we defined a network of side-chain and main-chain interactions between the bound ADP and the protein (Figure 6b, Table 3). This network includes lysine 191 (P-loop motif) and histidine 480 (MHD motif), as well as hydrophobic interactions from the GLPL motif, and other surrounding residues, resulting in a ligand-binding pocket with over 90% of the ADP surface area buried within the protein. The result of this extensive interaction surface is a binding pocket partially occluded by the positioning of the ARC2 subdomain, suggesting structural rearrangements will be required to allow nucleotide release (Figure 6c).

**Table 3.** Hydrogen bond interactions between the NRC1 NB-ARC domain and bound ADP, as defined by PDBe PISA (64).

| NRC1 residue | Atom | Ligand atom | Motif |
| --- | --- | --- | --- |
| Valine 158 | O | N<sub>6</sub> | hhGRExE |
|  | N | N<sub>1</sub> |  |
| Glycine 188 | N | O<sub>3</sub>B | GLPL |
| Glycine 190 | N | O<sub>2</sub>B, O<sub>3</sub>A | GLPL |
| Lysine191 | N | O<sub>2</sub>B | Walker A |
|  | Nζ | O<sub>2</sub>B |  |
| Threonine 192 | N | O<sub>1</sub>B | Walker A |
|  | Oγ1 | O<sub>1</sub>B |  |
| Threonine 193 | N | O<sub>1</sub>A | Walker A |
|  | Oγ1 | O<sub>1</sub>A |  |
| Arginine 326 | NH<sub>2</sub> | N<sub>3</sub> | RNBS-C |
| Histidine 480 | Nε2 | O<sub>3</sub>B | MHD |

Despite some regions of the refined electron density being well-resolved, the crystallographic R-factors for the final model remained high compared to other structures determined at similar resolutions. To help validate our model, we used the X-ray data set collected from SeMet-labelled protein crystals to generate an unbiased, experimentally-derived electron density map into which we docked the final model. We observed excellent correspondence of the docked model with this map (Figure 7). We were also able to position methionine residues at the experimentally determined heavy atom sites. Finally, electron density in this map confirmed the position of the bound ADP.

**Figure 7.**
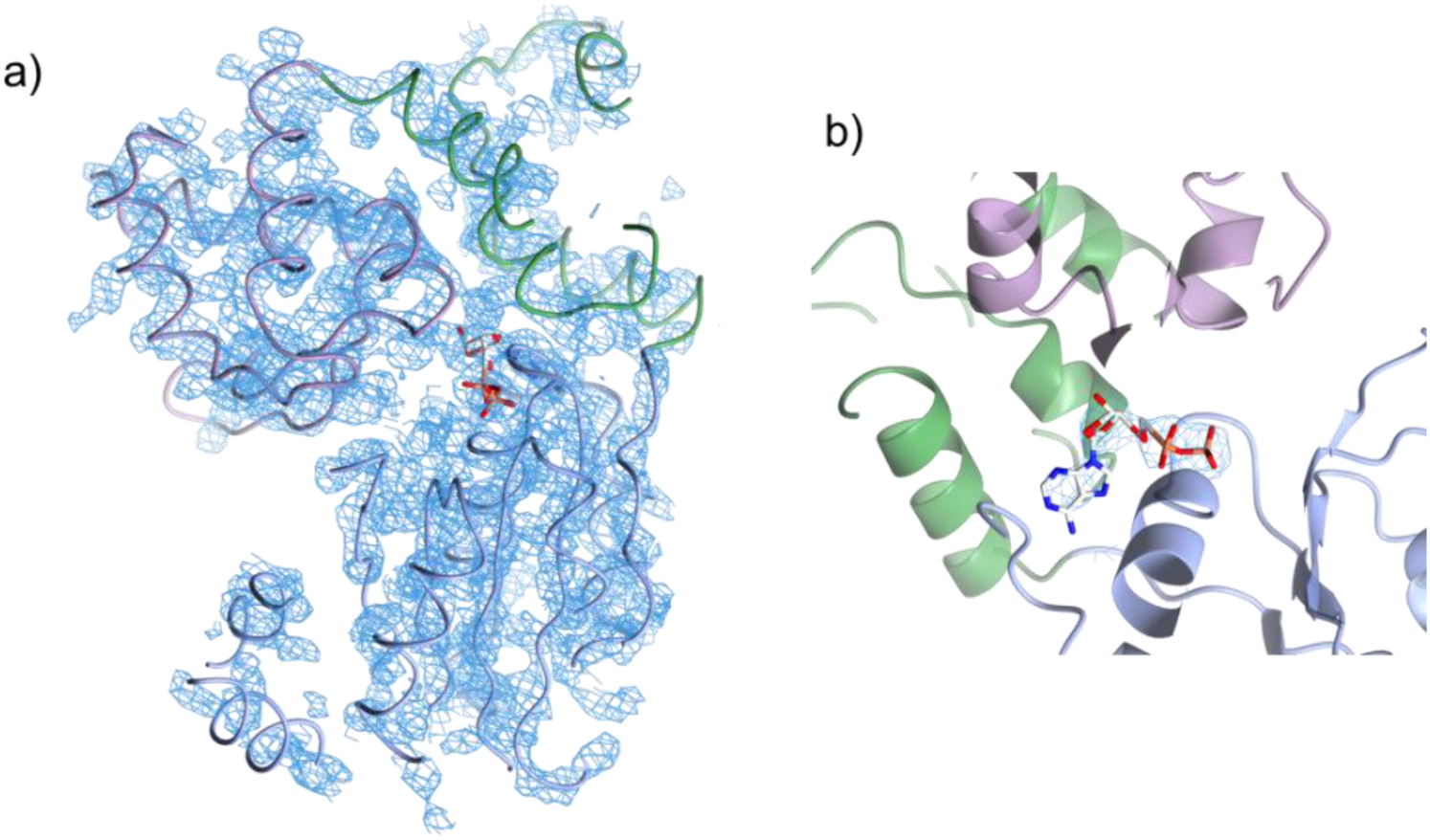
Experimentally-derived electron density map. a) Final model of the NRC1 NB-ARC domain, docked into the experimental electron density map. b) ADP docked into the experimental electron density map within the NB-ARC ligand binding site. The NB-ARC sub-domains are coloured as in Figure 5.

## Discussion

Plant NLRs are a key focus of research into plant-pathogen interactions due to their use in reducing pre-harvest crop losses to disease. Generalised models for plant NLR function and regulation have been proposed, primarily based on bioinformatic and *in vivo* studies (36, 37). Recent biochemical investigations have validated many of these predictions, and have developed a more mechanistic understanding of NLR regulation and activation (28, 39, 65, 66). Advances in our knowledge of NLR structures and their biochemical activities have relied on the limited number of NLRs where purified protein could be produced. The availability of new NLR proteins for in vitro study has generally led to novel insights into NLR behaviour (29, 67, 68).

The expression, purification, and initial biophysical characterisation of the NRC1 NB-ARC domain demonstrated it was suitable for further biochemical and structural experiments. Using this purified NRC1 NB-ARC domain we have confirmed the hypothesised structural homology between a plant and animal NB-ARC domains (85). The similarity of the NRC1 NB-ARC domain model to the equivalent region of APAF-1 validates the previous use of APAF-1 crystal structures to model and explain plant NLR mutant phenotypes (39,52,53).

Analysis of the nucleotide binding pocket of the NRC1 NB-ARC domain shows the presence of many residues from conserved NTPase motifs (Walker A, Walker B), as well as conserved NLR motifs that have established roles in protein regulation (GLPL, hhGRExE (53), and MHD). Further, the structure confirms that the NRC1 NB-ARC domain co-purifies with ADP suggesting that, like their animal counterparts, plant NLRs have high affinity for adenosine ligands and likely have either intrinsic ATPase activity, or have a binding preference for di-phosphate over tri-phosphate ligands. Low rates of ATP hydrolysis, and difficulties detecting ATPase activity, have previously been observed with studies of other NB-ARC proteins (40, 69, 70), and thus may constitute a regulatory mechanism for this family of proteins. As the NRC1 NBARC domain co-purifies with bound ADP, and not as apo-protein, it has not been possible to quantify ligand binding affinities. Most likely NB-ARC domains will be inherently unstable as apo-proteins, or will require additional factors (such as the presence of additional NLR domains, or external factors) to trigger ADP release.

The NRC1 NB-ARC domain structure in the vicinity of the bound ADP is unambiguously defined by the electron density, as are the relative orientations of the NB, ARC1 and ARC2 domains. However, due to the presence of other unresolved regions in the model, we suggest caution in interpreting additional details of the structure. Nevertheless, this limitation does not prevent analysis of the oligomerisation state of the protein in the crystal. In animal NLRs, NB-ARC domains form key interfaces with each other, contributing to the formation of inflammasomes (86, 87). Here, the NRC1 NB-ARC domain behaves as a monomer in solution and packing in the crystal lattice does not suggest a functionally relevant oligomerisation state. It is likely that other NLR domains, or structural rearrangements on activation, are necessary for NB-ARC domain oligomerisation.

We have identified methods for producing a soluble, stable plant NLR NB-ARC domain to allow in vitro investigation of its properties. This resource, and our data, demonstrates how structural and biochemical studies can improve our knowledge of NB-ARC domain functions. Ultimately, in vitro studies of full-length plant NLRs will be required to place these new findings in a more biological context, and give a more global understanding of plant NLR regulation.

## Materials and methods

### Construct design

Initial bioinformatics analysis to identify a likely NB-ARC domain was performed by submitting the NRC1 ORF to the Pfam server (50, 51). The extent of the LRR and coiled-coil domains were predicted using LRR Finder (54) and LRRsearch (55), and CCHMM (71) and MARCOILS (72). Prediction of secondary structure elements used PHYRE2 (56) and was supplemented with disorder predictions using RONN (57).

### Cloning

Wild-type NRC1 NB-ARC domain was amplified by PCR to include compatible ends for Infusion® cloning using forward primer: AAGTTCTGTTTCAGGGCCCGCCTGTGGTTGAGGAAGATGATGTGG and reverse primer: ATGGTCTAGAAAGCTTTACTCTGTCGTAGCCTCTTGCCAGC. The 1078 bp region was then cloned into pOPIN-S3C, which encodes a cleavable N-terminal 6xHis-SUMO tag, and pOPIN-F, which encodes a cleavable N-terminal 6xHis tag, as per manufacturer’s instructions (Clontech). Mutant sequences were synthesised by Genscript and cloned into pOPIN-S3C and pOPIN-F as above.

### Protein expression

Lemo21(DE3) (NEB) *E. coli* cells transformed with the NRC1 NB-ARC pOPIN-S3C plasmid were grown in LB media with 100 µg/mL carbenicillin at 37°C for 16 hrs with constant shaking. 20 mL of culture was used to inoculate one litre Powerbroth® (Molecular Dimensions). Cultures were grown at 37°C until an OD_600nm_0.5-0.7 was reached and then rapidly chilled on ice prior to induction of expression with 1mM isopropyl β-D-1-thiogalactopyranoside (IPTG). Following induction, cultures were grown at 18°C for 16 hrs, and harvested by centrifugation at 3500*g* for 10 mins. Pellets were then stored at −80°C until use.

Insect cell-derived pellets expressing pOPIN-F:NRC1 NB-ARC were produced by baculovirus-mediated transformation of *Sf9* cells by the Oxford Protein Production Facility and stored at −80°C until use.

### Protein purification, *E. coli*

Bacterial cell pellets were re-suspended in lysis buffer (50mM Tris-HCl pH 8.0, 50mM glycine, 5% glycerol, 500mM NaCl, 20mM imidazole, EDTA-free protease inhibitor cocktail (Roche)) and lysed by sonication. The extract was clarified by centrifugation at 37,000*g* for 30 mins at 4°C. Immobilised metal ion affinity chromatography (IMAC) was performed using HisTrap FF columns (GE Healthcare) pre-equilibrated with 50 mM Tris-HCl pH 8.0, 50 mM glycine, 5% glycerol, 500 mM NaCl and 20 mM imidazole. Single-step elution was performed by the addition of the same buffer supplemented with 500 mM imidazole. Eluate was then directly loaded onto a HiLoad Superdex 75 26/60 preparative gel filtration column, pre-equilibrated with A4 buffer (20 mM HEPES pH 7.5, 150 mM NaCl). 8 mL fractions were collected and assessed by SDS-PAGE.

### Protein purification, insect cells

Cell pellets were resuspended in 50mM Tris-HCl pH 7.5, 500 mM NaCl, 30 mM imidazole, 0.2% v/v TWEEN20 and EDTA-free protease inhibitor tablets and lysed using a cell disruptor (Constant Systems Ltd) at 14,000 psi. Following lysis, 10-50 μL Benzonase was added, and incubated at room temperature for 15 mins. Cell debris was removed by centrifugation at 30,000*g* for 30-40 mins at 4°C. The supernatant was passed through 0.45 μm filters (Sartorius), with additional incubation with Benzonase if required, before loading onto a 5 mL His-Trap FF column pre-equilibrated with equilibration buffer (50 mM Tris-HCl pH 7.5, 500 mM NaCl, and 30 mM imidazole). Bound proteins were eluted with equilibration buffer including 500 mM imidazole. The eluate was then loaded directly onto a Superdex S75 gel filtration column pre-equilibrated with 30 mM Tris-HCl pH 7.5, 200 mM NaCl, and 1 mM TCEP. Buffer was then exchanged for 50 mM HEPES pH 7.5 with 150 mM NaCl either by gel filtration following N-terminal tag cleavage, or by concentration and dilution.

### Generation of selenomethionine protein

*E. coli* carrying the pOPIN-S3C:NRC1 NB-ARC plasmid were grown overnight in minimal media consisting of M9 (0.6% w/v Na_2_HPO_4_, 0.3% w/v KH_2_PO_4_, 0.05% w/v NaCl, 0.1% w/v NH_4_Cl), 0.4% w/v glucose, 2 mM MgSO_4_, 0.1 mM CaCl_2_, 0.8% w/v each valine, phenylalanine, isoleucine, leucine, glutamate, lysine, arginine, serine, threonine, tyrosine, histidine, glutamine, tryptophan, 0.001% thiamine, supplemented with carbenicillin. Cells were pelleted by centrifugation at 1900*g* for 10 mins and washed twice in 20 mL M9 plus carbenicillin, before being re-suspended in 20 mL M9 with carbenicillin. This was used to inoculate 2 L of minimal media, and cultures incubated at 37°C for 6-8 hrs with constant shaking. When cells reached an OD_600_ of 0.3, 0.01% w/v threonine, lysine, phenylalanine, 0.005% w/v leucine, isoleucine, valine, and 0.006% w/v selenomethionine was added. Cells were grown for a further 45 mins at 37°C before induction overnight at 18°C, initiated by the addition of 1 mM IPTG.

### N-terminal tag removal

N-terminal 6xHis-SUMO (pOPIN-S3C) and N-terminal 6xHis (pOPIN-F) tags were removed by incubation with 3C protease overnight at 4°C. Protease and cleaved tag were then removed by subtractive IMAC prior to gel-filtration.

### Protein quantification

For crystallisation, analytical gel filtration, and CD, protein concentrations were determined using absorbance at 280 nm with a Nanodrop spectrophotometer. For ATPase assays and ligand quantification, concentrations were determined using infrared absorbance using a Direct Detect®.

### Crystallisation

Sitting-drop vapour-diffusion experiments identified suitable crystallisation conditions for native protein from the PACT screen (QIAGEN), with optimisation identifying the best conditions for crystal formation as 6 mg/mL protein combined 1:1 with 8% PEG 1500, 0.1 M MMT (MES, malic acid and Tris,) pH 4.5. Crystals typically formed in under 48 hrs. Selenomethionine-derivative protein was crystallised in 0.1 M MMT buffer pH 5.0 with 15% PEG 1500. Crystals were cryoprotected in mother liquor with the addition of 20% ethylene glycol.

### Data collection and processing

Data were collected at Diamond Lightsource (UK) beamlines I02 (native) and I04 (SeMet) under beamline proposal mx7641. Data were processed using the Xia2 pipeline (73) using XDS (74) (native data), DIALS (75) (SeMet), POINTLESS (76, 77) and AIMLESS. The structure was determined by SAD using the PHENIX AutoSol pipeline (78), and structure building was performed in COOT (79). Refinement was performed with Refmac5 (77, 80) and PHENIX, and validation performed with tools in PHENIX, COOT and MOLPROBITY (88).

### Analytical gel filtration

Analytical gel filtration was performed using a Superdex ™ S75 10/300 GL column equilibrated in 20 mM HEPES pH 7.5 and 150 mM NaCl. A standard curve was produced using a low-molecular weight calibration kit (GE Healthcare) for molecular weight estimation. Protein was diluted to 50 µM in equilibration buffer prior to loading 100 µL onto the column.

### Differential scanning fluorimetry

DSF was performed as described in Walden *et al*. 2014 (81). Briefly, assays used a Bio-Rad CFX96 ™ real-time thermal cycler with a thermal range from 20°C to 90°C, increasing by 1°C per minute. Fluorescence readings were taken every 0.2°C interval. Protein was prepared by dilution to 1 mg/mL in 20 mM sodium phosphate buffer pH 7.5, with 5 µL of dilution added to 45 µL reaction buffer (40 µL 20 mM sodium phosphate buffer pH 7.5 plus 5 µL 25x Sypro Orange (Invitrogen)). Assays were performed four times in 96-well plates, with five technical replicates per plate. Analysis was performed as described by Walden *et al*. 2014.

### Circular dichroism

Circular dichroism was performed using a Chirascan Plus instrument (Applied Photophysics). Protein was diluted in 20 mM sodium phosphate buffer pH 7.5 to a concentration of 0.1 mg/mL. Spectra were taken from 180 to 260 nm at 20°C in 0.5°C steps with four readings taken per step, averaged and buffer subtracted. Spectra at 20°C were taken from two independent purifications. Table 1 shows estimated secondary structure content from representative spectrum as analysed using CONTIN on the DichroWeb server (82, 83).

### Ligand quantification

ADP and ATP quantification assays were modified from Maekawa et al. (29) and Williams et al. (40). Bound nucleotide was released from the protein by denaturation followed by centrifugation (which pellets unfolded protein). The resulting supernatant was then divided equally and either directly incubated in the luciferase assay (to determine the ATP bound fraction), or first treated with pyruvate kinase to convert ADP to ATP prior to incubation (ADP bound fraction was determined by subtracting the ATP calculated in the untreated sample from that of the treated sample). Here, 15-20 µM protein was boiled in triplicate for 7 mins and centrifuged at 16000*g* for 12 mins at 4°C to precipitate denatured protein. The supernatant was removed and flash-frozen in liquid nitrogen until used. To convert released ADP to ATP for quantification, supernatant was diluted 1 in 10 in pyruvate kinase reaction buffer (85 mM Tris-HCl pH 7.4, 1.7 mM MgSO_4_3.5 mM phosphoenolpyruvate, 23U pyruvate kinase (Sigma-Aldrich)) for 30 mins at room temperature. To quantify co-purifying ATP, protein extract was treated as above, with pyruvate kinase replaced by an equal volume of 50% glycerol. Reactions were stopped by heating to 95°C for 7 mins and centrifuged at 16000*g* for 12 mins. 100 µL of supernatant was added to an equal volume of ATP bioluminescent assay buffer (Sigma Aldrich cat. FLAA) and photon emissions read immediately using a Clariostar plate reader (BMG). Each assay included an internal control of ADP, which underwent identical treatment to allow reference to a standard curve of ATP. Assays were performed three times and normalised by buffer subtraction and calibration with the internal standard. Data from all the experiments are presented as box plots, generated using R v3.4.3 (https://www.r-project.org/) and the graphic package ggplot2 (89). The centre line represents the median, the box top and bottom limits are the upper and lower quartiles, the whiskers are the 1.5× interquartile range and all of the data points are represented as dots.

## Acknowledgements

We thank Dr Simon Williams for advice performing ligand quantification assays, Dr Miriam Walden for advice performing DSF and CD analysis, and the Oxford Protein Production facility for expression screening and insect cell expression. We also thank Drs Adam Bentham and Marina Franceschetti for help preparing the manuscript. This work was supported by the Biotechnology and Biological Sciences Research Council (BBSRC, UK), including grant nos. BB/P012574, BB/J004553 and the John Innes Foundation.

